# A Multi-Crystal Method for Extracting Obscured Signal from Crystallographic Electron Density

**DOI:** 10.1101/073411

**Authors:** Nicholas M Pearce, Anthony R Bradley, Patrick Collins, Tobias Krojer, Radoslaw P Nowak, Romain Talon, Brian D Marsden, Sebastian Kelm, Jiye Shi, Charlotte M Deane, Frank von Delft

## Abstract

Macromolecular crystallography is relied on to reveal subtle atomic difference between samples (e.g. ligand binding); yet their detection and modelling is subjective and ambiguous density is experimentally common, since molecular states of interest are generally only fractionally present. The existing approach relies on careful modelling for maximally accurate maps to make contributions of the minor fractions visible (*1*); in practice, this is time-consuming and non-objective (*2*–*4*). Instead, our PanDDA method automatically reveals clear electron density for only the changed state, even from poor models and inaccurate maps, by subtracting a proportion of the confounding ground state, accurately estimated by averaging many ground state crystals. Changed states are objectively identifiable from statistical distributions of density values; arbitrarily large searches are thus automatable. The method is completely general, implying new best practice for all changed-state studies. Finally, we demonstrate the incompleteness of current atomic models, and the need for new multi-crystal deconvolution paradigms.

**One Sentence Summary:** Normally uninterpretable map regions are reliably modelled by deconvoluting superposed crystal states, even with poor starting models.

## Background

Besides its use for resolving the overall 3D structure of bio-molecules, macromolecular X-ray crystallography (MX) is deployed extensively to observe small changes to known structures, especially compound binding in ligand-discovery and -development projects. Arriving at the final model once initial electron density estimates are available (after “phasing”), relies on a long-established and rarely-questioned paradigm: cycling between building atoms into the current density estimate and computationally optimising the model against the measured data (“refinement”). The latter improves the calculated phases and yields more detailed density that should reveal additional model omissions and errors; the process is assumed to converge on a model that fully describes the crystal’s content.

In practice, convergence is never convincingly achieved. Much density both strong and weak invariably remains unexplained (“noisy”), hence the aphorism that “refinement […] is never finished, only abandoned” (*5*), and hence too the “R-factor gap” (*6*), which has obdurately resisted all methodology advances. More recent work has shown that conventional single-conformation models are too simplistic to describe the crystal (*7*–*9*); and that electron density features far weaker than the conventional cut-off reflect model deficiencies rather than measurement error (*10*, *11*).

Evidently then, near convergence, conventionally-calculated (sigmaA-weighted (*12*)) density derived from a single dataset is necessary but insufficient to complete the model, as it shows a superposition of states that is currently impossible to de-convolute algorithmically. Nearly-complete models with discrete yet uninterpretable superpositions are common in systematic studies of perturbations involving few atoms, such as ligand binding, photochemical changes or radiation damage. Since even strong biophysical effects are contingent on crystal packing or integrity, only a subset of the crystal may transition away from the ground state, often even after extensive optimization of the experiment. Finally, all current modelling approaches ultimately rely on shape-matching, and density superpositions are susceptible to interpretation errors and bias (such as the problem of the “Ligand of Desire” (*2*)).

Existing methods to auto-generate multi-conformer models (*8*, *9*) are not relevant when changes are chemical, and moreover have had little take-up, presumably because neither is explicit modelling involved nor have robust validation criteria emerged to allay long-cultivated fears of over-fitting (*13*). Approaches from time-resolved crystallography (*14*) apply only to specialised experiments.

## New Approach

In order to obtain unencumbered views of the changed, non-ground state, and extract the appropriate signal from conventional single-dataset density, we recast the problem as a multi-dataset, 3D background correction problem. An accurate estimate of the background can be obtained by averaging near-convergence density, in real space and after local alignment, from dozens (>30) of independently measured but approximately identical ground state crystals. Subtraction of a suitable fraction of this background estimate from the near-convergence density of a dataset containing a putative changed state yields a residual partial-difference map that we call an *event map* and that is in general fully interpretable (Figure 1):

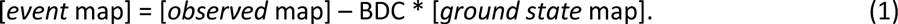

**Figure 1.**
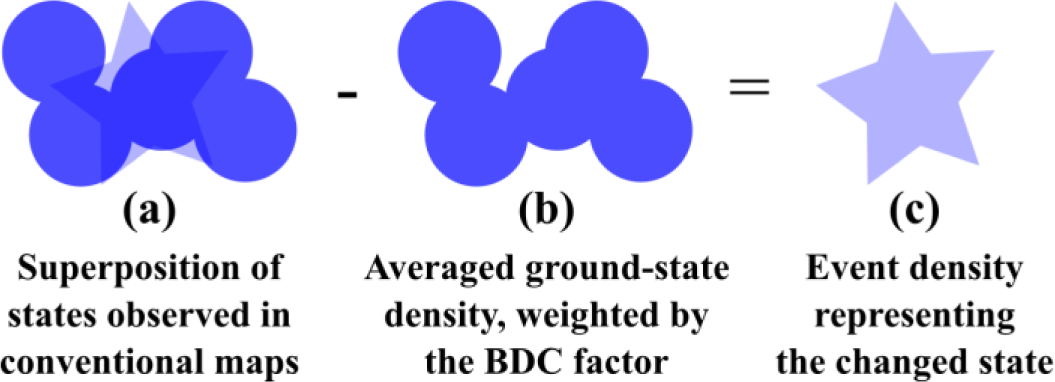
Schematic in 2D of how 3D background subtraction reveals changed-state density. With pixel intensity representing electron density strength, (a) shows the superposition of changed (20%) and ground state (80%) densities, while (b) shows the ground state density, estimated from the mean of ground-state measurements, and adjusted by applying a weighting (BDC=0.8). (c) The density that remains after subtracting background yields the best estimate of the changed state.

Our new method – Pan-Dataset Density Analysis (PanDDA) – comprises: the characterisation of a set of related crystallographic datasets of the same crystal form; the identification of (binding) events; and the subtraction of ground state density to reveal clear density for events. Identifying the optimal Background Density Correction factor (BDC) is essential for extracting the best signal, illustrated schematically in Figure 2.

**Figure 2.**
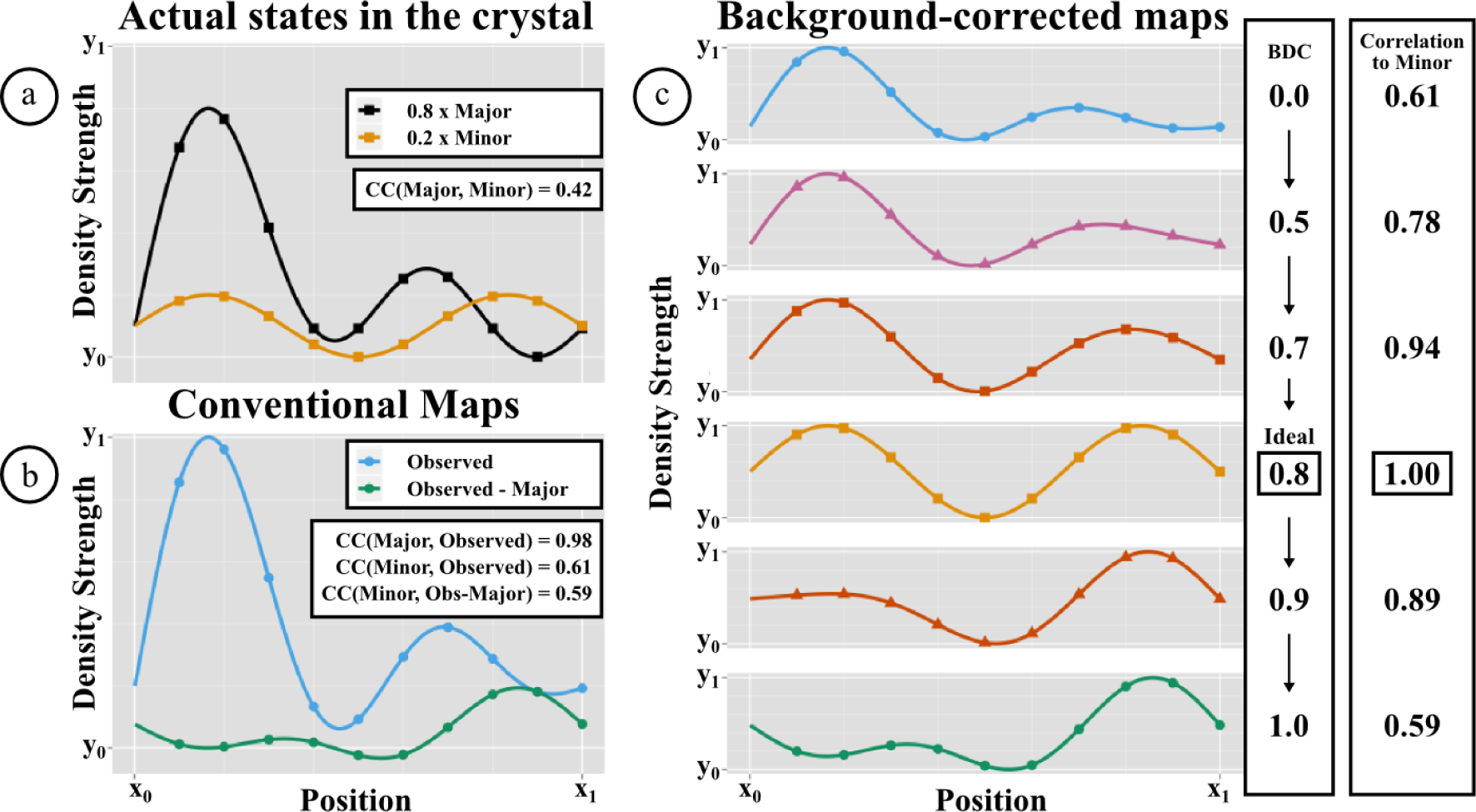
Minor conformations are obscured in conventional maps, but revealed by background correction. 1D simulations are used to illustrate 3D electron density. (a) The actual crystal contains 80% major (black) and 20% minor (orange) states, which are largely dissimilar (correlation: 0.42). (b) Conventional (2mFo-DFc) maps (blue) show only the superposition, which resembles the major far more than the minor state (correlations: 0.98 and 0.59; in practice, the scale is arbitrary). Isomorphous difference (Fo-Fo) maps (green) show the subtraction of the full-occupancy major state from the observed dataset, and only resemble the minor state where the major state has low density (right side). (c) “Event maps” (scaled for comparison), generated as in equation (1) for different values of BDC, reveal the minor state optimally for one value of BDC (0.8). BDC=0.0 corresponds to the observed density, and BDC=1.0 to a F_o_-F_o_ map.

The method builds on the principle of isomorphous difference (F_o_-F_o_) maps (*15*), but analyses many maps simultaneously by (a) locally aligning maps in real space to bypass the requirement of strict isomorphism, and (b) directly comparing the best estimate of true electron density, namely sigmaA-weighted (2mF_o_-DF_c_) maps from late-stage refinement, ensuring maps are correctly scaled.

Using multiple maps allows a Z-score measure to be calculated, reflecting how significantly each dataset deviates from the ensemble of datasets at each point in space. Z-scores are assembled into spatial *Z-maps*, where clusters of large Z-scores are an objective and statistically meaningful measure for potentially interesting crystallographic signal – *events* – such as a binding ligand. Using Z-maps addresses the common pitfall of over-interpreting density that is in fact ground state density, since in such cases, Z-scores will be small. Equally importantly, Z-maps also make it possible to identify weak changed states (e.g. weakly-bound ligands) that do not yield strong difference (mF_o_-DF_c_) density.

Finally, the precise localisation of each change enables reliable background subtraction at that site, because BDC can be estimated as the value for which the ground state-subtracted map is locally least correlated to the ground-state map, relative to a normalising global correlation across the unit cell (Supplementary D). Using the average map both reduces noise of the ground-state estimate and thereby of the event map, and provides a less-biased estimate of the true ground-state, which a single-dataset map cannot, as it is inherently biased by the model. A correct estimate of BDC results in event map density for only the changed configuration of the site, including protein backbone and side-chain conformations induced by the change.

## Results

Crystallographic fragment screening (*16*, *17*) represents an extreme case of changed-state studies, because it attempts to observe in electron density the rare and often low occupancy binding events that occur when a relatively large (200-1000) library of weak-binding “fragment” compounds (150-300Da, 100μM-10mM) (*18*, *19*) are added individually or as cocktails to a series of equivalent crystals. Conventionally, the analysis is challenging as it involves inspecting a lot of 3D space – the whole unit cell in all datasets – for convincing evidence of bound fragments (“hits”). In contrast, PanDDA directly eliminates the thousands of strong electron density blobs with no statistical significance, objectively identifying only regions that are unique to each dataset; the ground state datasets are provided by the many hit-free crystals.

Applied to a series of fragment screens (Table 1), PanDDA yielded markedly more hits than manual inspection of density, far more quickly and all with high confidence (Figure 3 & Figure 4; Supplementary Figure S1-Figure S4), in both known binding sites and new allosteric sites (Figure 4d). Several fragments induced significant reordering of sections of the protein that could only be modelled with PanDDA event maps (Figure 4a-c, Figure S1a-c), whilst also enabling the identification of mislabelled ligands and the discovery of experiment errors (Figure S1d-f, Figure S2d-f). Models erroneously built into misleading conventional density could be discarded with statistical confidence, and the binding of chemically elaborated hit compounds could be analysed more reliably. Full experimental details and complete descriptions are provided in Supplementary A. The method also effectively disambiguates density in conventional ligand-binding studies with ligands co-crystallised and a sub-optimal number of ground-state datasets (Supplementary B).

**Figure 3.**
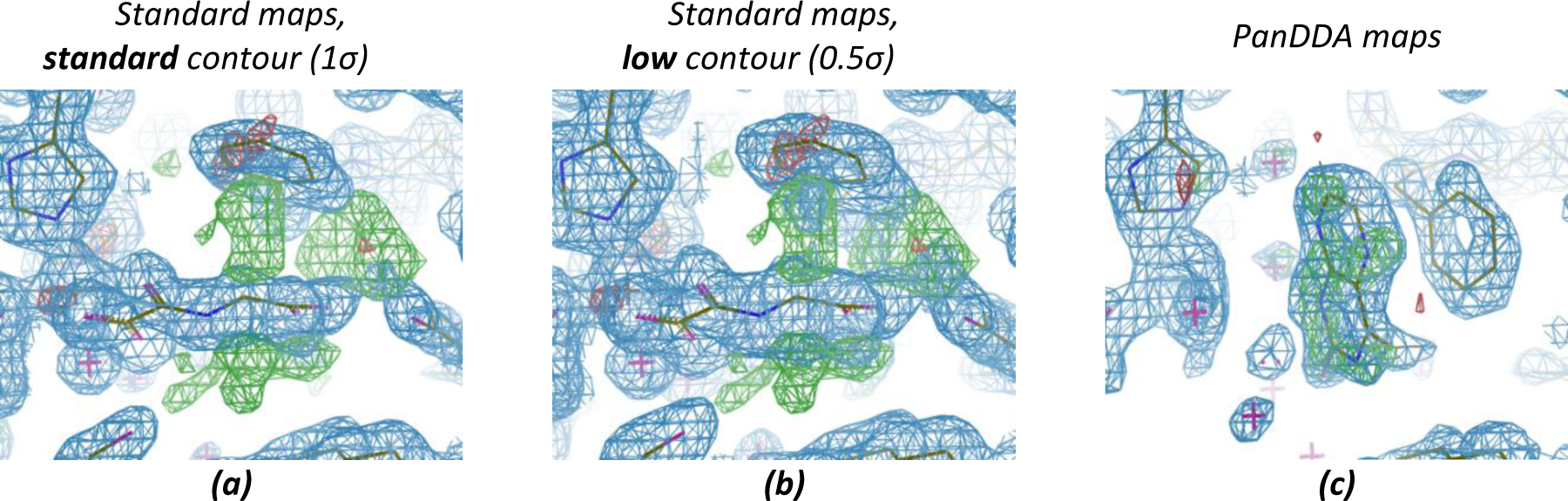
PanDDA maps clearly show detail obscured by conventional maps. JMJD2D fragment screening dataset x401 at 1.48Å. (a,b) Conventional maps (2mFo-DFc, blue, contour as indicated; mFo-DFc, green/red, ±3σ) are dominated by the NOG co-factor analogue bound in the majority fraction of the crystal, whereas (c) the event map (blue, 2σ, BDC=0.9) and the Z-map (green/red, ±4) unambiguously reveal both ligand and associated changes in protein conformations.

**Table 1.** Hit rates from fragment screens before and after use of PanDDA. All fragment screens consisted of a single soaked compound per dataset. An identified site comprises more than 2 binding ligands that are not heavily interacting with crystal contacts. Number of hits was determined as number of datasets containing a bound ligand. Hit rate was calculated as percentage of datasets containing bound ligands.

| Protein | JMJD2D | BAZ2B | SP100 | BRD1 |
| --- | --- | --- | --- | --- |
| Datasets | 226 | 200 | 116 | 292 |
| Resolution Range (Å) | 1.1-2.6 | 1.5-2.5 | 1.3-2.7 | 1.5-3.6 |
| Identified Hits (Human / PanDDA) | 2 / 24 | 3 / 9 | 0 / 2 | 29 / 40 |
| Identified Hit Rate (%) (Human / PanDDA) | 0.9 / 10.6 | 1.5 / 4.5 | 0 / 1.7 | 9.9 / 13.7 |
| Identified Sites (Human / PanDDA) | 1 / 5 | 1 / 1 | 0 / 1 | 1 / 2 |

**Figure 4.**
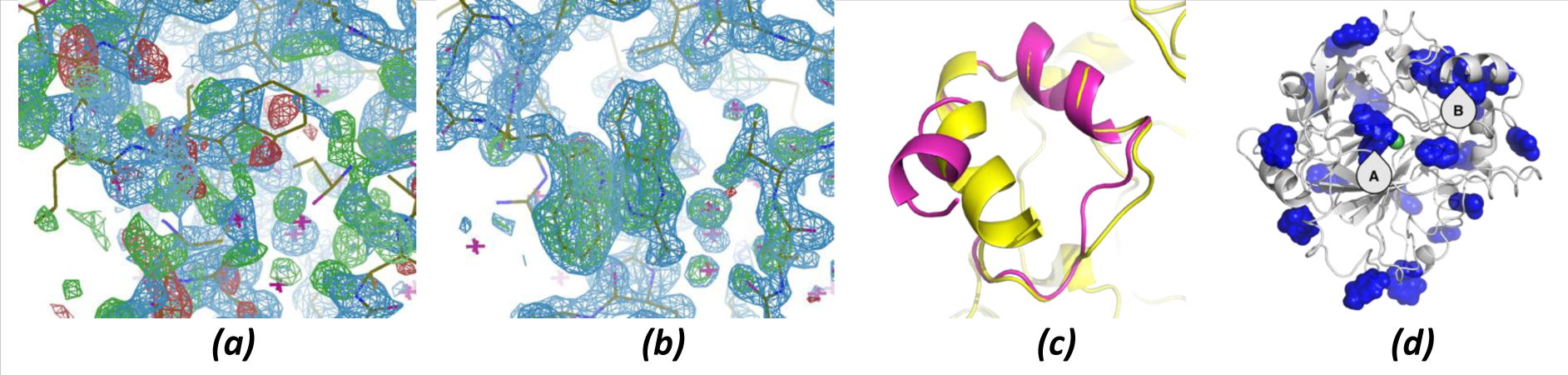
PanDDA maps reveal complex minor conformations and identify allosteric binders. In JMJD2D dataset x402 (1.45Å), (a) conventional maps (contoured as in Figure 3a) show a complex superposition difficult to model using the reference conformation (shown), while (b) in PanDDA maps (contoured as in Figure 3c) it can be modelled easily. (c) Final models for the unbound (yellow) and bound (magenta) conformations show the large conformational change. (d) Fragments are detected to bind all over the surface of JMJD2D, revealing potential allosteric sites, including the peptide-binding groove (site A) and the large helix reordering (site B).

Strikingly, detection of weak binding events is simple even when phases are far from convergence (Figure 5).

**Figure 5.**
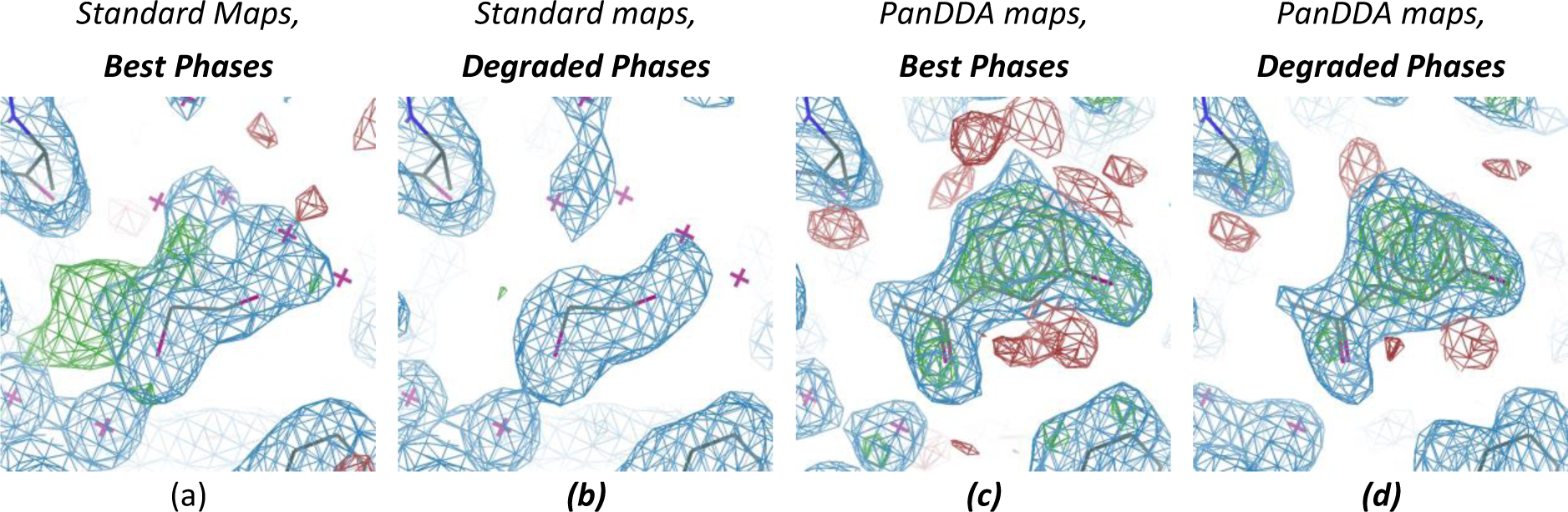
Weak ligand identification remains straightforward when phases are degraded. BAZ2B datasets were re-analysed using a deliberately sabotaged reference model, introducing a ∼30° phase error and increasing Rwork and Rfree by ∼12% for all datasets. Shown here is the weak hit (occupancy: 0.64) in dataset x492, contoured for different maps as labelled: (a,b) 1.78Å 2mFo-DFc (blue, 1σ) and mFo-DFc (green/red, ±3σ). (c,d) 1.79Å event (blue, 2σ) and Z-maps (green/red, ±3). Rwork/Rfree are 0.18/0.21 and 0.30/0.32 for best and degraded phases respectively. BDCs for best and degraded phases are 0.77 and 0.73 respectively, and although the quality of the density for the ligand is reduced, ligand identification is no more difficult.

## Validation

Model validation is a long-established bedrock of crystallographic analysis (*13*), and crucially requires a model that is numerically stable in refinement. To enable this, we generate an atomic ensemble model that reflects the crystal content by combining the ground state with the changed state modelled from event maps, with initial occupancies of 2*(1-BDC), as discussed in Supplementary D. These models are indeed well-behaved. However, we discovered that many, some built into strong event density, would be considered invalid (Figure S6) by the subjective but best-practice criterion (*2*) of visual assessment of agreement between model and conventional OMIT maps.

To address this, we formulated the following strong objective validation principles:

1. The changed-state partial model must conform to calculable numerical criteria (Table S2). We adopt established requirements: a high correlation between the model and the observed density (RSCC>0.7); that the model must not move under refinement (low rmsd before *vs* after); and that ligand B-factors must be comparable to those of surrounding residues. We also apply a new metric, that modelling and refinement should result in negligible difference density around the site (RSZD<3) (*20*). These metrics are fully defined in Methods and shown for all models in Table S3-Table S6.
2. The ground state partial model is considered an immutable component of the crystal, with a status similar to common restraints (*e.g.* geometry or non-crystallographic symmetry), as in general there is not enough diffraction information to propose otherwise. Thus, the ground state model needs to be fully complete before incorporation into the ensemble, and during further cycles of model building, it may not be altered. To stabilise refinement, it may need to be strongly restrained to the original ground state model (by external restraints using *e.g.* PROSMART (*21*)).
3. The primary event density must always be available when disseminating such models.

We note that the infrastructure for criterion 3 does not currently exist in the PDB (*22*); and refinement programs do not yet support some external restraints that we predict will be important for numerical stability at low resolution or for very low occupancy at high resolution, in particular restraining relative B-factors to stabilise the occupancy. Both are the subject of future work.

In general, only the changed state will be of primary scientific interest in the refined model, with the ground state essentially an experimental artefact. Unlike the artefacts inherent in any crystal structure, here they are explicitly declared and need not be inferred by further analysis. Structure repositories, whether public (PDB) or internal, would ideally support this by removing the ground state for normal 155 use.

## Discussion

The PanDDA algorithm improves on current methods not only with dramatically better signal-to-noise, but also by providing rigorous measures of confidence. This allows far more subtle changes to be modelled, whose importance will be experiment- and context-dependent: in ligand development, evidence of weak binding is now known to be productive for optimising binding potency (*23*).

We thus propose a new standard practice for ligand binding and other changed-state studies, namely collecting a series of ground state datasets before proceeding with the putative changed-state datasets, to provide the contrast necessary to see the changes of interest. Approximately 30 datasets are required for *full* convergence of the statistical model (Supplementary I), an experiment that can be completed within hours at modern synchrotron beamlines with fast pixel detectors (*24*) and sample automation (*25*), and that needs to be performed only once per crystal form. To address the other bottleneck, the logistics of analysing large numbers of datasets, the PanDDA implementation includes graphical tools and various command-line options. However, the method also works when fewer than 30 datasets are available (Supplementary B), the trade-off being potentially reduced quality of the event maps; determining the break-even number of datasets for a given case is the subject of future work.

The PanDDA method is applicable and effective at any resolution, though at lower resolutions, as maps become less precise, higher occupancies of changed states will in general be required for them to be detected by Z-score. What matters most is the consistency of ground-state models so that they can be well-represented by an average; therefore, in regions of crystals that tend to vary stochastically, such as crystal contacts, statistical confidence is reduced similarly to low resolutions.

As the algorithm is a contrast-maximisation approach, event map density for changes appears somewhat stronger than density for unchanged atoms (typically, surrounding protein). In practice, this is not problematic, as unchanged conformations do not require modelling anyway, as more fully discussed in Supplementary D.

In principle, the approach will allow comparisons between different crystal forms of the same protein. However, since functionally important conformational changes are not only common in such cases but by their nature affect the functionally interesting regions, algorithmic treatment of the local alignment is complex and the topic of future work.

Our results upend a long-held tenet in macromolecular crystallographic model building, that to visualise subtle features requires optimal phase estimates and thus a model as complete and *globally* error-free as possible (*1*). Conscientiously observed, this places a heavy time burden on the analysing scientist as it demands multiple iterations of modelling for each dataset. The PanDDA approach makes this both practically and theoretically unnecessary: a single local modelling step fully validates an interpretation, even when the model retains problems elsewhere.

More generally, we submit that a qualitative shift in approaches to generating crystallographic models is now due. PanDDA addresses one class of experiments, those involving induced local changes, but all problems of uninterpretable density, and indeed some of the R-factor gap (*6*), should be addressable by analogous map deconvolution methods. Multi-dataset experiments are no longer difficult; nevertheless, existing tools focus on pursuing a single, representative dataset through averaging (*26*). Instead, what will be key is establishing methods for targeted perturbations of poorly ordered regions, along with rigorous algorithms for reconstructing and visualising discrete states, and for subsequent model validation.

## Acknowledgements

The authors thank Randy Read and Garib Murshudov for productive conversations, and Luis Ospina for discussions regarding the statistical model. All data were collected at Diamond Light Source beamline I04-1 as part of the SGC-Diamond I04-1 XChem partnership.

## Implementation

PanDDA is implemented in Python and relies heavily on the CCTBX (*27*). It has been tested extensively for robustness and usability by users of Diamond’s XChem fragment screening facility. Source code is available on bitbucket (http://bitbucket.org/pandda/pandda) or as part of CCP4 (*28*). A manual and tutorial are available at http://pandda.bitbucket.org. Processing 200-500 datasets on a 3.7GHz Quad-Core Intel Xeon with 32GB of RAM takes 3-10+ hours but runtime depends greatly on resolution binning and size of crystallographic unit cell.

## Data Availability

All crystallographic data, models, Z-maps and event maps are available at Zenodo (http://zenodo.org), with the following DOIs: BAZ2B: 10.5281/zenodo.48768; BRD1: 10.5281/zenodo.48769; JMJD2D: 10.5281/zenodo.48770; SP100: 10.5281/zenodo.48771. Models were built for those ligands that could be uniquely identified in the event density, except for those that interact extensively with the crystal contacts and are therefore unlikely to be biochemically relevant. Models have not yet been deposited in the PDB in order to ensure adherence to the essential validation principle 3 discussed above.

## Materials and Methods

An overview of the PanDDA algorithm is schematically outlined in Supplementary E.

### Dataset Preparation

The input to PanDDA is a series of refined crystallographic datasets, each consisting of a refined structure and associated diffraction data, including 2mF _o_−DF_c_ structure factors. These can come from any refinement program, as long as all datasets are refined using the same initial atomic model and the same protocol. All models of the protein must be identical, up to the numbering and labelling of atoms. All datasets used in this paper were prepared using the Dimple pipeline (part of CCP4 (*28*)), from reference models including solvent molecules; there is no requirement to remove solvent atoms from known binding sites.

### Structure and Map Alignment

To allow map voxels to be compared between crystals that are not exactly isomorphous, maps are aligned using the refined models as reference points.

The input protein structures are aligned using a flexible alignment algorithm (Supplementary F). Sections of the protein are aligned separately, to give alignment matrices for that section. The alignments generated from the structures are stored and are used to transform and thereby align the electron density maps.

### Handling Variations of Map Resolutions

To allow map voxels to be compared between crystals, maps have to be calculated at the same level of detail, even though crystals can diffract to a wide range of resolutions. For analysing a specific dataset, its full resolution is used; but for contributing to the analysis of a different dataset, higher resolution datasets are truncated to the resolution of the target dataset, while lower resolution datasets are ignored. Therefore, we analyse the collection of datasets at a number of resolutions, and high resolution datasets are used multiple times for characterisation at lower resolutions, but will only be analysed once, at their highest possible resolution. Maps are recalculated using truncated diffraction data at each different resolution limit. Thus, if processing in resolution bins of 1.0Å, 1.5Å, 2Å, and 2.5Å, a 1.2Å dataset would be analysed at 1.5Å, but also be used to build distributions at 2Å and 2.5Å.

Fourier terms omitted in a given map, as happens when reflections are unobserved and then effectively set to zero, lead to systematic changes in electron density throughout the unit cell that strongly affect the outlier analysis; strong low-resolution terms are particularly problematic. Therefore, reflections in all datasets are truncated to the set of miller indices common to all datasets; and for map calculation, all missing Fourier terms are estimated as DF_c_, which refinement programs perform automatically as long as the indices are correctly included in the reflection files.

Truncated 2mF_o_ −DF_c_ structure factors are Fourier-transformed to generate maps. These maps are aligned using the alignment transformations from the local alignment (Supplementary F).

### Statistical Model

Once maps for a particular resolution have been aligned, a statistical model is parameterised using the electron density of the *ground state* datasets. The aligned maps are placed on an isotropic Cartesian grid, and the electron density is sampled at each grid point of each dataset. The model treats the observed value of the electron density in dataset *i*, at grid point *m*, as being sampled from a distribution

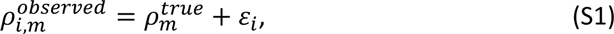

where 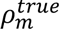 models the natural variation in the electron density at point *m*, independent of dataset, and *ε_i_* represents the experimental uncertainty in the electron density in dataset *i*. The variability of the 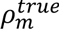 term accounts for the fact that the crystals are not identical, and that small local fluctuations may exist between the crystals. These areas are most likely to be in the crystal contacts, or flexible areas of the protein. 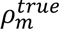 represents the “true” (unmeasurable) electron density for this crystal form, of which each crystal (and associated dataset) is a sample.

The simplest model is to assume that both the uncertainty in electron density values as well as variation in electron density at a point arising from differences between the crystals, can be modelled by a normal distribution. Therefore, if

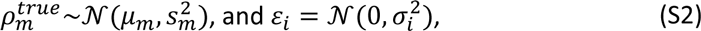

then

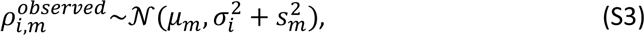

where *μ_m_* is the mean value of the electron density at point *m*, *s_m_* is the variance of the “true” electron density at point *m*, and *σ_i_* is the uncertainty in dataset *i*. Under this model, the parameters *μ_m_* are estimated by taking the un-weighted average of all of the ground state densities.

The mean ground state map is used to estimate the dataset uncertainty, *σ_i_*, for all datasets as follows. Subtracting the mean map from each dataset map we obtain a mean-difference map. By assuming that the experimental and model uncertainty in the electron density map are the major contributors to deviations from the mean map, the histogram of the mean-difference map values is used to estimate the total uncertainty of the dataset. Calculating the quantiles of a theoretical normal distribution 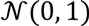 and plotting them against the quantiles from the mean-difference map, yields a Q-Q plot where the slope of the central portion of the map (between the ±1.5 theoretical quantiles) gives an estimate of the uncertainty of the dataset (Figure S11a). This is equivalent to the method used in Tickle (2012) for calculating the uncertainty of an electron density map (*20*).

To estimate *s_m_*, a maximum likelihood method is applied on our model in (S3), using the observed values 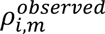, as well as estimates for *σ_i_* and *μ_m_* for the *ground state* datasets (Supplementary H). An example comparison of the ‘raw’ standard deviations of the grid points (simple standard deviation of electron density values, not accounting for observation error) and the ‘adjusted’ values is shown in Figure S12. This adjustment results in the majority of points having no variation that is not accounted for by the dataset uncertainties; the remaining points have non-negligible variation, with non-zero *s_m_*, and these indicate naturally variable regions.

### Calculation of Z-Maps

The parameterised statistical model allows the identification of areas of individual dataset maps that deviate significantly from the mean map: “events”. Z-scores are calculated by

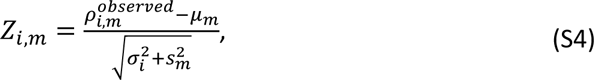

where large Z-scores indicate significant deviations from the mean map. The distributions of Z-scores for a particular dataset have improved normality compared to the simple differences from the mean (Figure S11b), as expected.

Regions of individual datasets are identified as significant by contouring Z-maps at Z=2.5, and filtering remaining blobs by a minimum peak value of Z=3 and a minimum volume of 10Å^3^ (volume of a water molecule is ∼30Å^3^). Neighbouring blobs are grouped together if the minimum distance between them is less than 5Å. These parameters were identified on the BAZ2B dataset, and found appropriate in subsequent studies and are therefore the current program defaults.

### Calculation of Event Maps

For identified events, the background density correction (BDC) factor is estimated as follows. Different fractions of the mean map are subtracted from the dataset map, and the correlation between the resulting map and the mean map is calculated both globally and for the area around the event, defined by the blob identified in the Z-map expanded by 1Å.

Globally, the dataset map looks similar to the mean map, so plotting the global correlation against the subtracted fraction yields a signal-to-noise curve, dropping off at a speed related to the noise in the dataset (green dashed line, Figure S7). Locally to the identified site, however, the dataset map is a superposition between something similar to the mean map and something that is unrelated (e.g. density of bound ligand). As more of the mean map is subtracted, the local correlation between the mean map and the resulting map (black dashed line, Figure S7) will decrease faster than the global correlation. Subtracting the local correlation curve from the global correlation curve, BDC is estimated where the difference between these two correlation curves is maximised (blue solid line, Figure S7). The final event map is calculated as in equation (1).

### Model Building and Refinement

Interesting sites are identified by Z-maps and modelling is performed using a combination of Z-maps and event maps, similarly to the way that mF_o_-DF_c_ maps may be used to guide the modelling of 2mF_o_-DF_c_ maps. Modelling takes place in the aligned *reference* frame, as defined in Supplementary F.

After modelling of the changed state, the new conformations of the protein are merged with the ground state model. Atoms in the ground state that are not present or have moved in the changed state are assigned to a previously unused conformer (e.g. C). Similarly, atoms in the changed state model that are not present in the ground state, or have moved, are assigned another unused conformer (e.g. D). Atoms that are not changed between the two states remain unaltered. The resulting ensemble models are then back-transformed, using the local alignments, to the original crystallographic frame, for refinement.

The models in Table 1 have then been refined as an ensemble using phenix.refine (*29*, *30*), under conventional resolution-dependant refinement protocols, with constrained occupancy groups corresponding to the bound and unbound structures to ensure that the occupancies of the bound and unbound states sum to unity.

Because of the methodical way in which the ensembles are generated, the changed state model can be extracted simply by removing the atoms corresponding to the changed ground state atoms (i.e. conformer C in the above example).

### Validation

The atomic model of the changed state is validated by 4 quality metrics (Table S2). Two are electron density scores, generated by *EDSTATS* (*20*): <u>RSCC</u> reflects the fit of the atoms to the experimental density, and should typically be greater than 0.7; while <u>RSZD</u> measures the amount of difference density that is found around these atoms, and should be below 3. The <u>B-factor</u> ratio measures the consistency of the model with surrounding protein, and is calculated from the B-factors of respectively the changed atoms and all side-chain atoms within 4Å. Large values (>3) reflect poor evidence for the model, and intermediate values (1.5+) indicate errors in refinement or modelling; for weakly-binding ligands, systematically large ratios may be justifiable. <u>RMSD</u> compares the positions of all atoms built into event density, with their positions after final refinement, and should be below 1Å.

## List of Supplementary Materials

Supplementary Figures S1-S13

Supplementary Tables S1-S7

Supplementary A - Fragment Screening Datasets

Supplementary B - Ligand Screening Studies

Supplementary C - Ligand Validation

Supplementary D - Background Density Correction

Supplementary E - PanDDA Implementation

Supplementary F - Flexible Alignment

Supplementary G - Uncertainty and Z-Map Calculation

Supplementary H - Estimation of Density Variability

Supplementary I - Statistical Model Convergence

